# Mapping resistance-associated anthelmintic interactions in the model nematode *Caenorhabditis elegans*

**DOI:** 10.1101/2023.04.26.538424

**Authors:** Elena G. Rehborg, Nicolas J Wheeler, Mostafa Zamanian

## Abstract

Parasitic nematodes infect billions of people and are mainly controlled by anthelmintic mass drug administration (MDA). While there are growing efforts to better understand mechanisms of anthelmintic resistance in human and animal populations, it is unclear how resistance mechanisms that alter susceptibility to one drug affect the interactions and efficacy of drugs used in combination. Mutations that alter drug permeability across primary nematode barriers have been identified as potential resistance mechanisms using the model nematode *Caenorhabditis elegans*. We leveraged high-throughput assays in this model system to measure altered anthelmintic susceptibility in response to genetic perturbations of potential cuticular, amphidial, and alimentary routes of drug entry. Mutations in genes associated with these tissue barriers differentially altered susceptibility to the major anthelmintic classes (macrocyclic lactones, benzimidazoles, and nicotinic acetylcholine receptor agonists) as measured by animal development. We investigated two-way anthelmintic interactions across *C. elegans* genetic backgrounds that confer resistance or hypersensitivity to one or more drugs. We observe that genetic perturbations that alter susceptibility to a single drug can shift the drug interaction landscape and lead to the appearance of novel synergistic and antagonistic interactions. This work establishes a framework for investigating combinatorial therapies in model nematodes that can potentially be translated to amenable parasite species.

## Introduction

Diseases caused by parasitic nematodes infect over one billion people and cause morbidity that reinforce poverty and profoundly increase years lived with disability. The clinical and subclinical impacts of parasitic nematodes are also responsible for significant losses in livestock production and companion animal health. Parasite treatment and control efforts in both human and veterinary medicine rely primarily on three drug classes: benzimidazoles, macrocyclic lactones, and nicotinic acetylcholine channel agonists. Although generally effective in prominent human helminth control and elimination campaigns [1,2], approved drugs exhibit suboptimal activity against some helminths, and anthelmintic resistance is a potential concern with the expansion of mass drug administration. Resistance to the major anthelmintic classes is already widespread in livestock [3] and a growing concern in small animals [4,5].

Approaches to mitigate resistance may include combinatorial treatments that improve efficacy against a specific helminth. Anthelmintics administered in combination are recommended or being considered for the management of lymphatic filariasis [6,7], whipworm [8,9], and strongyloidiasis [10]. In the veterinary realm, combination anthelmintics are used to expand the spectrum of antiparasitic activity and to help delay or overcome single-drug resistance [11–13]. While there are active efforts to better understand mechanisms of anthelmintic resistance in human and animal populations [14–19], it is unclear how resistance mechanisms that alter susceptibility to one drug affect the interactions and efficacy of drugs used in combination.

Validated resistance mechanisms in parasitic nematodes are restricted to mutations in the cytoskeletal targets of the benzimidazoles [5], but genetic tools in the model nematode *Caenorhabditis elegans* have helped to identify anthelmintic resistance mechanisms beyond drug target mutations [20]. These include mutations that affect the ability of drugs to accumulate within the worm by altering drug uptake, distribution, efflux, or metabolism [14,16,21,22]. Genetic mapping [18,23,24] and phenotypic observations [16,25] of anthelmintic responses in parasitic nematodes suggest that these resistance mechanisms are field relevant. It is also likely that non-target associated resistance mechanisms that affect the entry and movement of drugs within the worm have a greater potential to confer partial resistance or resistance across multiple anthelmintic classes.

Anthelmintics can be absorbed by nematodes via crossing the cuticle, diffusing through the cilia of the amphid neurons, or being ingested through the pharynx and intestine [26]. Mutations that alter these putative drug interfaces can selectively modulate anthelmintic activity in model nematodes [14,15,22,27–32]. We set out to investigate how genetic perturbations that impact drug entry and resistance to a given anthelmintic can alter the landscape of interactions between anthelmintics belonging to different classes. Using high-throughput phenotyping approaches in *C. elegans*, we mapped changes in the interaction landscape across the three primary anthelmintic drug classes and a selection of strains with mutations affecting putative routes of drug entry.

## Methods

### Code and data availability

All analytical code and raw tabular data can be found at https://github.com/zamanianlab/AnthelminticInteractions-ms. wrmXpress is publicly available at https://github.com/zamanianlab/wrmXpress. Raw imaging data is available upon request.

### Nematode strains and maintenance

*Caenorhabditis elegans* wild type (N2) and mutant strains were maintained on 6 cm plates seeded with *Escherichia coli* OP50 bacteria following standard protocols [33]. Six mutant strains were acquired from the *Caenorhabditis* Genetics Center (CGC); *bus-5(br19), agmo-1(e3016), nhr-8(ok186), eat-2(ad465), dyf-2(m160)*, and *che-1(p672)*. The selected strains carry mutations in tissues serving as putative interfaces for drug entry. *nhr-8(ok186) and eat-2(ad465)* have altered digestive system function [32], *bus-5(br19)* and *agmo-1(e3016)* have reduced cuticle integrity [28,29], and *dyf-2(m160)* and *che-1(p672)* have developmental defects in amphid neuron cilia [34].

### Drug treatment and development assay

Each drug was tested individually and in combination against all seven strains. Assay-ready plates (ARPs) were prepared using an automatic multichannel pipette (Eppendorf) to add 1 μL of 100X drug stock to each well in 96 well plates. ARPs were stored at -20°C until needed (<90 days).

On day 0, starved *C. elegans* plates were chunked to fresh 10 cm NGM plates seeded with *E. coli* OP50 and incubated at 20°C for 72 hours (or 96 hours for *eat-2(ad465)* due to a decreased growth rate). Strains were then bleach-synchronized and resulting embryos were titered to 3 embryos per μL in K media. Embryos for all but *bus-5(dc19)* were incubated in K media between 16-20 hours at room temperature on a nutator (Fisher S06622) at 13 rpm. *bus-5(dc19)* cannot be hatched in polypropylene tubes due to extreme adherence and was alternatively hatched in glass tubes that were shaken at 180 rpm or on unseeded NGM plates. These alternative hatching methods led to similar drug responses (**S2 Fig**). After 16-20 hours of hatching, resulting L1s were re-titered to 1 embryo per μL in K media. ARPs containing 100x drug stock were equilibrated to room temperature and 50 μL L1 worms (50 worms total) were added with 50 μL HKM (concentrated *E. coli* HB101, K media, and kanamycin (final concentration 25 μg/mL)) to individual wells. Plates were sealed with breathable film (Diversified Biotech BERM-2000) and incubated for 48 hours in a humid chamber at 20°C with shaking at 180 rpm.

After 48 hours, plates were washed with an AquaMax 2000 (Molecular Devices) in preparation for imaging. Liquid was aspirated at a probe height of 5 mm, leaving 100 μL liquid in each well. The plate was then shaken on the “fast” setting for 30 seconds and 280 μL of M9 or 1.4% 1P2P was dispensed, after which the worms soaked for 6 min to allow all worms to settle to the bottom of the well. Liquid was again aspirated at a probe height of 5 mm and 280 μL of either 140 mM sodium azide or M9 was dispensed to fill the well. After washing, each well was imaged at 2X using an ImageXpress Nano (Molecular Devices).

### Phenotypic measurements

Image data was analyzed as previously described [35]. Briefly, a custom Cell Profiler pipeline was implemented using wrmXpress [35], which segmented worms and extracted various morphological features from computationally straightened worms. MajorAxisLength (the length of each worm) was used for all downstream analysis. Outliers were defined and pruned as observations that fall outside the IQR by at least 1.5xIQR. The mean length was calculated for each well within an assay plate, with each well representing a single technical replicate of a drug-dose-strain combination.

### Data analysis and statistics

All data was analyzed with the R statistical software and publicly available packages, including tidyverse, drc, and tidymodels [36–38]. All data were normalized by dividing the mean length from each treatment well by the mean length of the control population from the corresponding plate (1% DMSO). For single-drug dose response experiments, a four-parameter log-logistic model was fit to the mean of normalized lengths from each well. Curves and EC_50_ values were calculated and plotted for each individual replicate and as a whole. Robustness of inferences were checked and confirmed across other normalization schemes (**S1 Fig**). Standard errors were calculated for the EC_50_ of each strain-drug combination. Briefly, the standard error of the log(EC_50_) was calculated over replicates and the defined interval around the geometric mean of the log(EC_50_) was converted from the log to linear scale. The relative potencies of each drug were compared between mutant strains and the wild-type (N2) strain using the EDcomp() function of the drc package to derive p-values.

For two-drug isobolograms, worm lengths were normalized by dividing the mean length from each treatment well by the mean length of the control population (1% DMSO) from the corresponding biological replicate. SynergyFinder 2.0 [39] was used to assess drug interactions, using percent inhibition (relative to mean length of untreated animals) as the response value and the zero interaction potency (ZIP) model [40]. Heat maps were generated for normalized lengths, then inhibition values were calculated by taking (1 - normalized value x 100) to be used in ZIP synergy score calculations. Antagonism was defined by ZIP synergy values less than -10 and synergy was defined by ZIP synergy values greater than 10.

## Results

### High-throughput measurement of anthelmintic drug effects on *C. elegans* development

Our goal was to investigate anthelmintic drug interactions across *C. elegans* genetic backgrounds that are hypothesized to differentially alter drug entry. As a first step, we optimized a high-throughput imaging assay to measure the effects of individual drugs on worm development in the wild type strain N2. Bleach-synchronized L1 stage animals were seeded into liquid culture microtiter plates and worm lengths were quantified after 48 hours of incubation in the drug. 8-point dose response curves were generated for albendazole sulfoxide (AZS), ivermectin (IVM), and levamisole (LEV) (**Fig 1**). Each assay included three wells per drug condition (technical replicates) and at least three independent assays were carried out across both a common drug stock and independently generated drug stocks. Assay variation was mostly associated with drug stock preparation and we moved forward with a common drug stock for all biological and technical replicates in subsequent assays and focused our inferences on relative measures of drug modulation. The calculated confidence intervals for N2 EC_50_ values fall into ranges similar to previous investigations [17,31,41].

**Fig 1.**
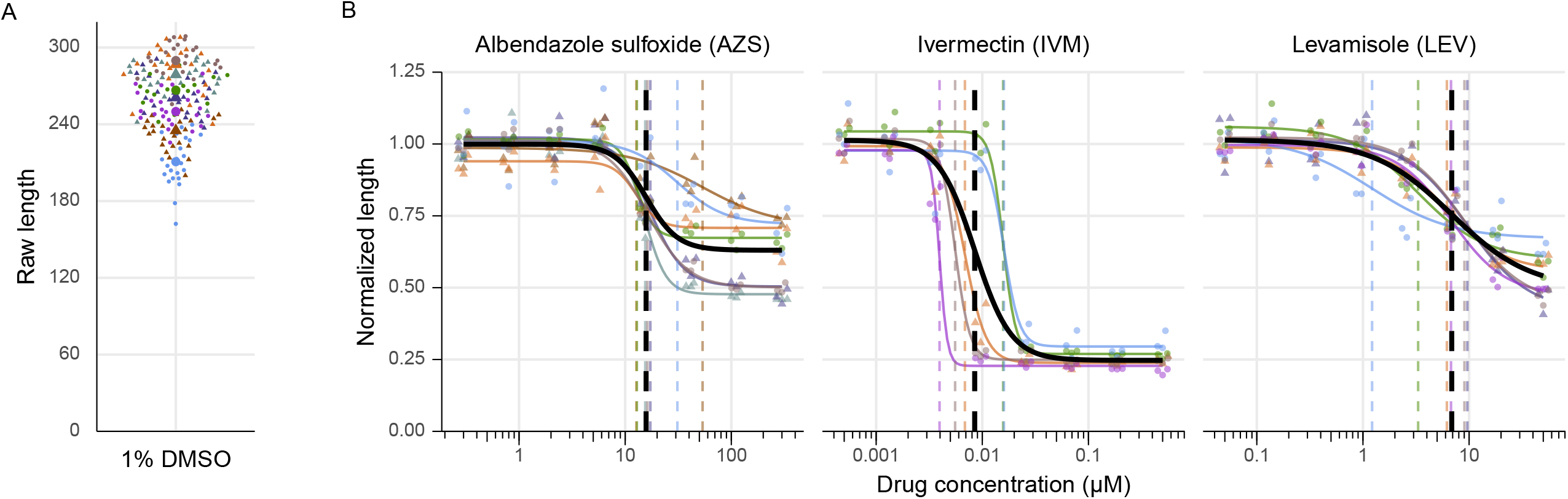
Dose responses of wild type *C. elegans* development for three primary anthelmintics. **A)** Dot plot showing the distribution of raw length in control worms (1% DMSO), colored by replicate. Triangles represent replicates performed with individually prepared drug stocks, while circles represent replicates performed with aliquots of a shared drug stock. **B)** Dose response curves of three anthelmintic drugs, albendazole sulfoxide (benzimidazole), ivermectin (macrocyclic lactone), and levamisole (nicotinic acetylcholine channel agonist) in wild type *C. elegans* (N2). Dose response curves (solid lines) and EC_50_ values (dashed vertical lines) are grouped and colored by replicate. The global fit was calculated using all data and is depicted in black. EC_50_ means and associated standard errors can be found in **Table 1**.

### *C. elegans* strains with mutations in putative drug entry pathways show variation in anthelmintic response

We next measured drug responses across a panel of six mutant strains with genetic perturbations in one of three putative routes of drug entry: the digestive tract (*eat-2(ad465)* and *nhr-8(ok186)*), the amphid (*che-1(p672)* and *dyf-2(m160)*), and the cuticle (*agmo-1(e3016)* and *bus-5(br19)*). We performed 8-point dose response experiments to derive EC_50_ values for comparison with drug responses in wild type (N2) worms (**Fig 2, Table 1**). The digestive tract mutant *nhr-8(ok186)* displays a two-fold increase in sensitivity to albendazole sulfoxide (p < 0.0001), while *eat-2(ad453)* displayed increased sensitivity to levamisole (p < 0.0001). Neither digestive mutant background resulted in significant changes in animal growth in response to ivermectin. The cuticle mutant *agmo-1(e3016)* showed a near two-fold increase in sensitivity to albendazole sulfoxide (p = 0.001) and levamisole (p = 0.002), but does not alter ivermectin sensitivity in our assay. The cuticle mutant *bus-5(br19)* did not shift the potency of any of the three tested drugs compared to wild type worms. Both amphid mutants displayed an increase in resistance to ivermectin (p < 0.0001), and *dyf-2(m160)* showed a slight increase in resistance to albendazole sulfoxide (p = 0.049).

**Fig 2.**
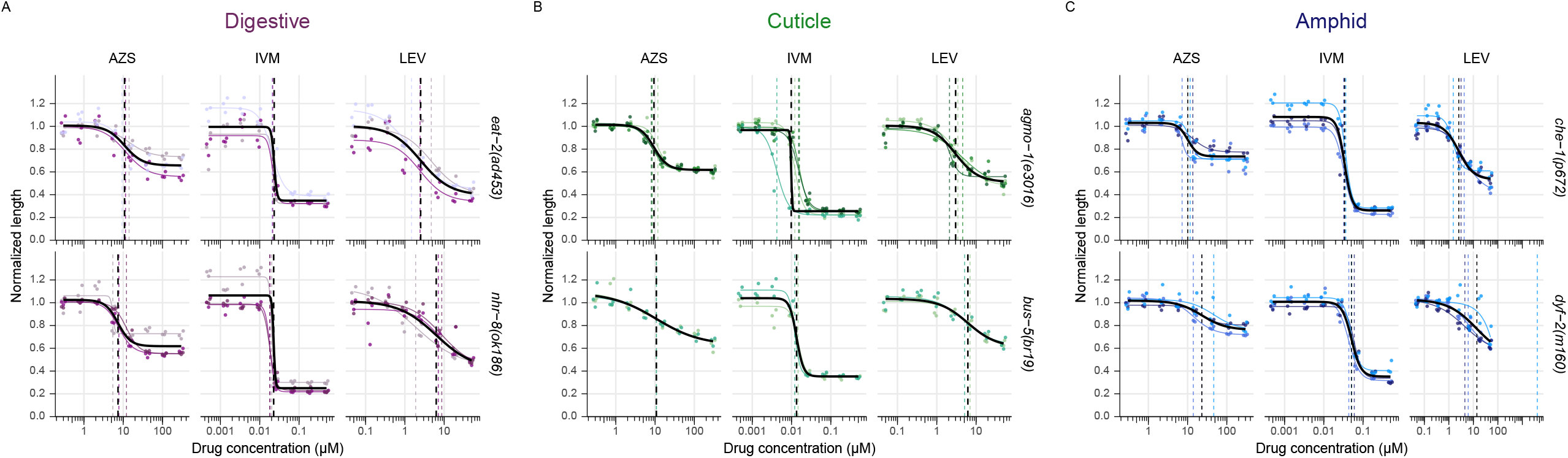
Anthelmintic dose response curves for strains with mutations in the digestive tract, cuticle, or amphid. Vertical dashed lines represent EC_50_ values. Colored curves and dashed lines represent individual biological replicates and the bold black line depicts the overall curve and its associated EC_50_. EC_50_ values and confidence intervals are reported in **Table 1. A)** *eat-2(ad453)* displayed an increase in sensitivity to levamisole (p < 0.0001) and *nhr-8(ok186)* displayed an increase in sensitivity to albendazole sulfoxide (p < 0.0001). **B)** *agmo-1(e3016)* displayed an increase in sensitivity to albendazole sulfoxide (p = 0.0013) and levamisole (p = 0.0022), while *bus-5(br19)* showed no significant change in response compared to wild type (N2) worms. **C)** Both amphid mutants show increased resistance to ivermectin (p < 0.0001) and *dyf-2(m160)* also showed a modest increase in resistance to albendazole sulfoxide (p = 0.0494).

Our assay identified significant differences in relative drug potency associated with mutations in putative drug entry pathways but did not recapitulate some previous findings. For example, *eat-2(ad465)* and *nhr-8(ok186)* alter susceptibility to ivermectin [42, 31]. This may be a function of different phenotyping schemes and endpoints, as assays differ in their sensitivity and ability to measure subtle drug effects across strains with variable growth rates and in different liquid or solid plate culture conditions. Armed with at least one mutant strain within each drug entry pathway that modulates a response to at least one tested drug, we next tested how these mutant strains affected interactions across the primary drug classes.

### Anthelmintic interactions are altered by resistance-associated mutation

The effects of pairwise combinations of the three primary drugs (AZS, LEV, and IVM) on worm development were measured across wild type and mutant strains and reported using zero interaction potency (ZIP) synergy scores [40]. Assays were set up using an 8-point isobologram approach with the same culture and environmental conditions as the dose response experiments. Concentration ranges were chosen considering the EC_50_ values calculated in the wild type (N2) dose responses. The effects of the pairwise drug interactions on worm development were calculated across these concentration ranges for wild type and mutant strains **(Fig 3)**. This phenotypic response landscape was used to generate ZIP scores and map the synergy landscape across the three drug interactions (**Fig 4**). ZIP values close to zero indicate drug responses that were not significantly different than the expected additive effects of the two drugs. Drug synergy was defined by ZIP synergy scores greater than 10 and drug antagonism was defined by ZIP synergy less than -10 [39].

**Fig 3.**
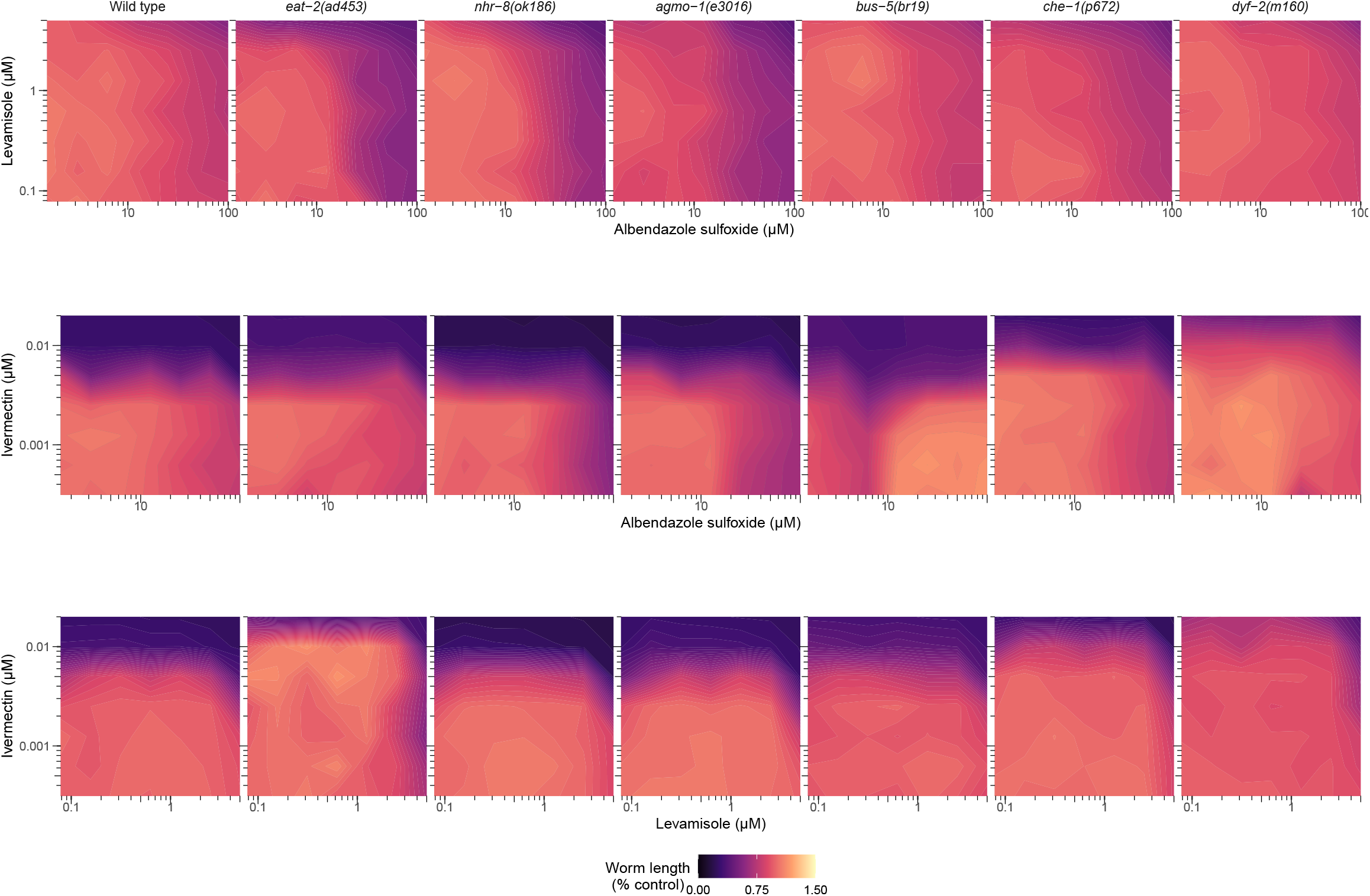
Contour plots displaying the effects of anthelmintic drug combinations on worm development in wild type and mutant strains. Drug responses are plotted across N2 and six mutant strains exposed to combinations of LEV and AZS **(A)**, IVM and AZS **(B)**, and IVM and LEV **(C)** across selected concentration ranges.

**Fig 4.**
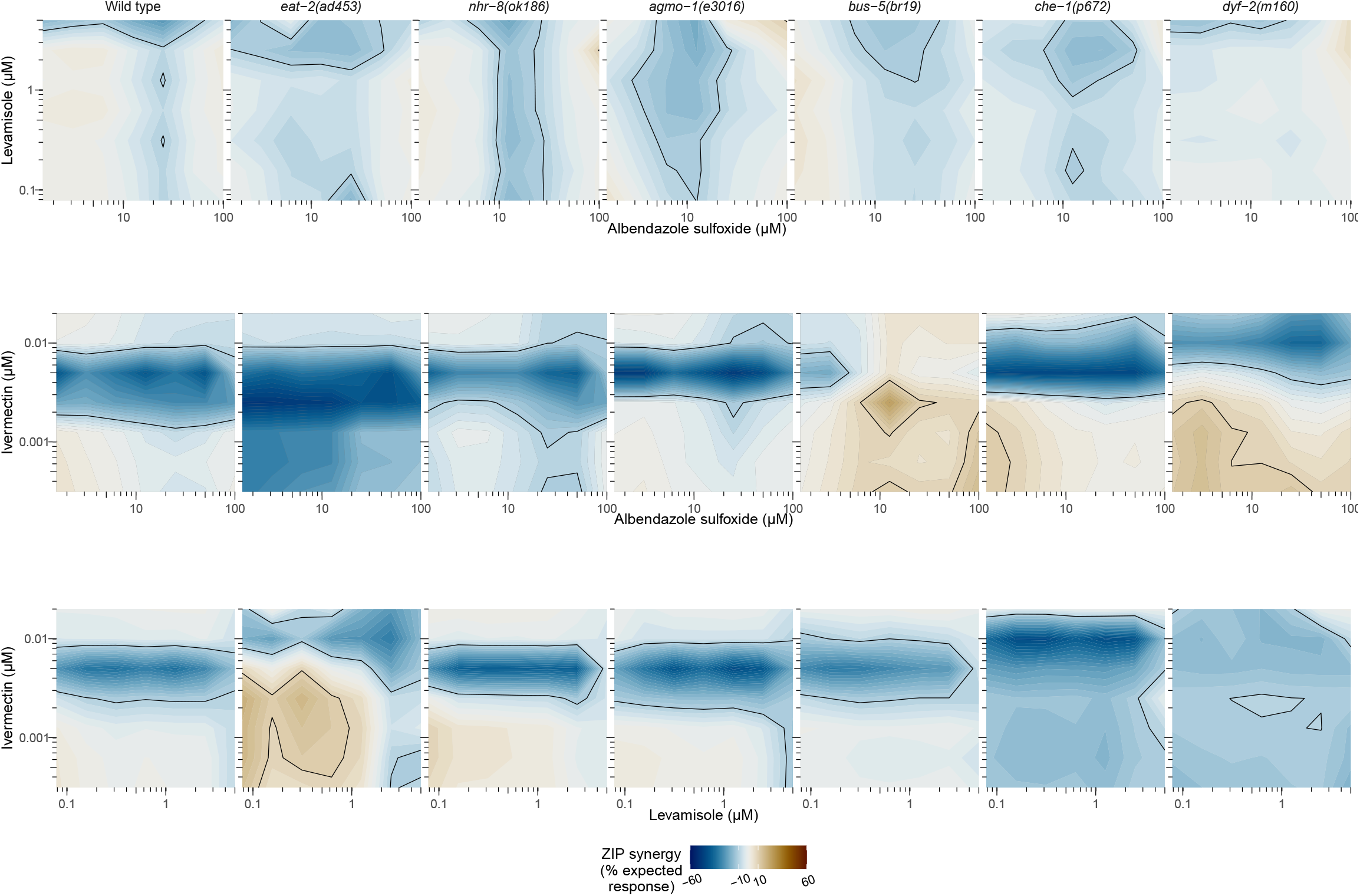
Contour plots displaying the synergy landscapes of anthelmintic drug combinations in wild type and mutant strains. Drug synergy values (ZIP scores) are plotted across N2 and six mutant strains exposed to combinations of LEV and AZS **(A)**, IVM and AZS **(B)**, and IVM and LEV **(C)** across selected concentration ranges. Regions of synergy (ZIP > 10; brown) and antagonism (ZIP < -10; blue) are contoured with a black line.

The different mutant backgrounds altered the wild-type drug interactions in both modest and significant ways. Interactions between LEV and AZS are primarily additive across wild type and mutant strains, with minor modulation of antagonism at high concentrations of LEV (> 2.5 μM) and middle range concentrations of AZS (6.25 - 25 μM). More pronounced shifts in drug interactions were observed for the drug pairings that included ivermectin. The wild type profiles for the IVM-AZS and IVM-LEV interactions are very similar, with a band of antagonism centered around 5 nM IVM. Different genetic perturbations lead to ZIP landscape shifts that include the movement of this baseline band of antagonism and the appearance of synergistic interactions.

In the IVM-AZS interaction, the digestive mutant *eat-2(ad453)* displays antagonism at lower concentrations of IVM (< 0.01 μM) compared with wild type. Interestingly, the cuticle mutant *bus-5(br19)* displays a synergistic interaction centered around 2.5 nM IVM and 12.5 μM AZS, as well as at lower concentrations of IVM (< 2.5 nM) paired with high concentrations of AZS (> 50 μM). The movement of the baseline antagonism of amphid mutant *dyf-2(m160)* to a higher concentration of IVM might reflect the independent effect of this mutant on IVM susceptibility. This mutant also displays synergistic effects at low concentrations of both IVM (< 2.5 nM) and AZS (< 50 μM).

The IVM-LEV interaction also displays some unique shifts across genetic perturbations. The most significant changes occur in the *eat-2(ad453)* and *dyf-2(m160)* backgrounds. There is an upward shift in the concentration of IVM associated with antagonism in the *eat-2(ad453)* strain compared with wild type worms, as well as the appearance of synergistic effects between IVM and LEV at lower concentrations of both. In the *dyf-2(m160)* strain, there is a diffusion of the narrow band of baseline IVM-LEV antagonism across the entire range of tested concentrations.

## Discussion

Overall, we see a number of significant changes to the anthelmintic interaction landscape in strains carrying mutations associated with putative drug entry pathways. Most but not all of the observed shifts occur in mutant strains that independently affect responses to a single drug in the tested interaction. Antagonistic drug interactions were present across all three drug pairings in the wild type background, although more pronounced in the interactions that included IVM. This antagonism is centered around a pharmacologically relevant concentration (5 nM) and may reflect a form of single-agent dominance [43] whereby the effects of IVM on development are realized earlier than either AZS or LEV. IVM at higher concentrations is likely saturating the developmental delay phenotype due to its faster onset.

In general, we expect antagonistic and synergistic drug effects to be explained by a complex mixture of factors that require much deeper dissection of both pharmacodynamic and pharmacokinetic interactions within the worm. Beyond timing of drug effects, different anthelmintics act on targets that are found in both distinct and overlapping tissues and cell populations. For example, the targets of AZS and IVM have overlapping expression in neuronal cells [44], which could contribute to the antagonism we see between these two drugs at higher IVM concentrations. We observed that *dyf-2* introduces synergy at lower concentrations of IVM and AZS. The decreased potency of IVM in this strain might allow for AZS effects to be realized before saturation of the developmental inhibition phenotype by IVM. Similarly, the *eat-2* strain introduces synergy at lower concentrations of IVM and LEV, which could be explained by the reciprocal pattern where the increased potency of LEV allows it to exert its developmental effects more quickly with respect to IVM. Our data also reveals that novel interactions can arise in mutants that do not respond differently to the individual drugs in the interaction pair. While *bus-5* did not show significantly different responses to IVM and AZS alone, it displayed an altered IVM-AZS interaction landscape compared with N2.

The concentrations of drug used in this study fall within the range of previous *C. elegans* studies and may provide some insight into pharmacologically relevant drug exposure in parasitic nematodes [46]. While the translation of these *in vitro* studies is unclear, it has been observed that higher concentrations of drug are required to elicit effects in *C. elegans* as a likely function of lower cuticle permeability [47]. The maximum plasma concentration (C_max_) has been investigated in patients administered combinations of IVM, AZS, and LEV [48] and these concentrations fall within (IVM and LEV) or near (AZS) the concentration ranges we investigated in the isobolograms.

Additional studies are needed to better understand the principles underlying both antagonistic and synergistic anthelmintic interactions across different genetic backgrounds. Here, we focused on a small number of genetic perturbations associated with drug entry pathways and observed some drastic shifts in the nature of drug interactions. Investigating the impacts of mutations that alter drug transport and metabolism would provide a more complete picture of how the efficacy of combinatorial therapies is altered by resistance mechanisms. It may be possible to carry out similar studies in parasitic nematodes with abundantly accessible life stages amenable to genetic perturbation [49,50]. We expect that expansion of this line of work will lead to a better understanding of pharmacological considerations that are often ignored as it relates to nematodes.

## Acknowledgments

Some strains of *C. elegans* were provided by the Caenorhabditis Genetics Center, which is funded by the NIH Office of Research Infrastructure Programs (P40 OD010440). We thank members of the Zamanian laboratory for their insightful comments on this manuscript.

---

**S1 Fig**. N2 dose-response data depicted using varying normalization schemes. Analysis was performed considering four normalization schemes, and patterns and curves of best fit vary minimally between methods. **A)** Phenotypic data was normalized by dividing individual values by the average of the control (1% DMSO). **B)** Max-min normalization using the highest concentration of drug as the minimum and the DMSO control as the maximum. **C)** The normalization procedure in (A) preceded by a square root transformation. **D)** The normalization procedure in (B) preceded by a square root transformation.

**S2 Fig**. Two alternative egg hatching methods used to generate L1-synchronized *bus-5* populations show comparable drug responses. Dose response data is shown for eggs hatched on unseeded NGM plates (black) and in glass tubes (blue).

