## Supplementary figures and images for "Mapping resistance-associated anthelmintic interactions in the model nematode *Caenorhabditis elegans*"

### S1 Fig

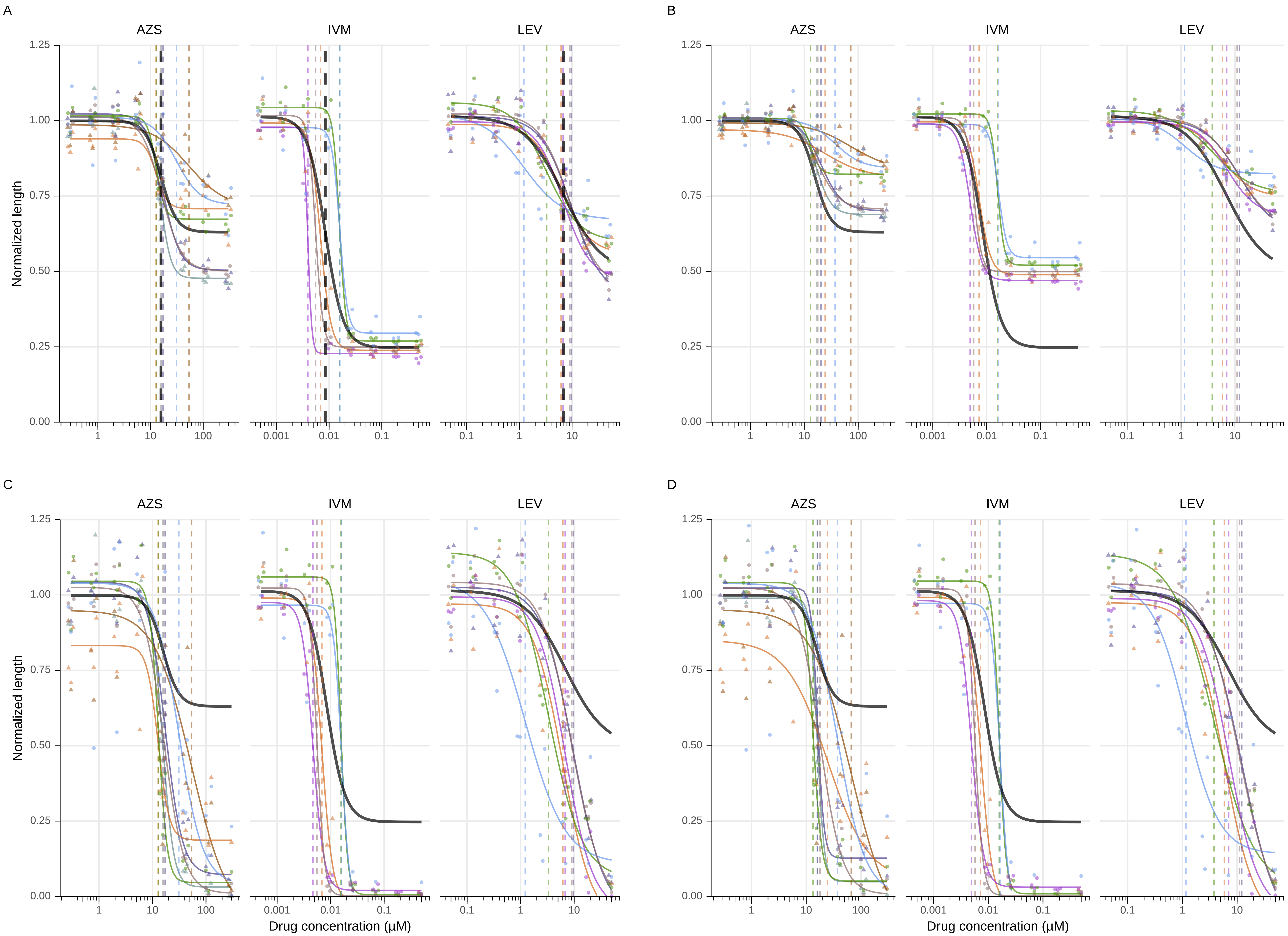

### S2 Fig

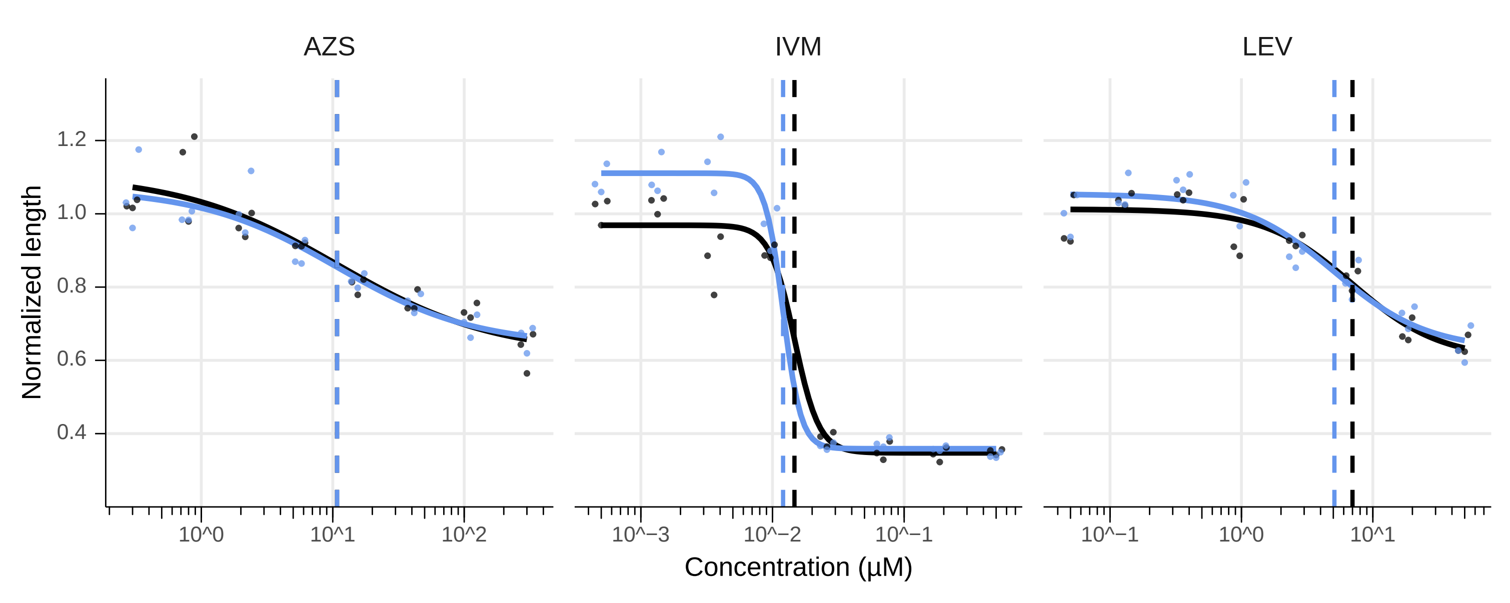
